## Supporting information for "Automatic structure-based NMR methyl resonance assignment in large proteins"

### Contents

**Figure S1.** Parametrization of the distance cutoff $d_{\mathrm{cut}}$ and the probability of expected methyl-methyl NOE contacts, $p_{\mathrm{NOE}}$.

**Figure S2.** Accuracy of the methyl assignments obtained for the different values of NOE probabilities over a range of NOE distance cutoffs, $d_{\mathrm{cut}}$.

**Figure S3.** Determination of the optimal number of individual assignment runs for automatic methyl resonance assignment with MethylFLYA

**Table S1.** Methyl resonance assignments by MethylFLYA, MAGMA, MAP-XSII, and FLAMEnGO2.0.

**Figure S4.** Sources of errors in the automatic methyl resonance assignments generated by MethylFLYA.

**Table S2.** Summary of errors in the automatic methyl resonance assignments generated with MethylFLYA

**Figure S5.** Parameter optimization for automatic NOESY peak picking with CYPICK.

**Figure S6.** 2D $^{13}$C($\omega_1$)-$^{13}$C($\omega_2$) projections of 3D CCH NOESY (ATCase, HSP90) or 4D HCCH NOESY spectra (EIN).

**Table S3.** MethylFLYA computation times (h) for different combinations of input NMR data, as in Figure 3.

**Table S4.** Results of the CYPICK application to the 3D CCH NOESY (ATCase, HSP90) and 4D HCCH NOESY (EIN) spectra.

**Figure S7.** Intersection of assignments generated with different automatic methyl assignment protocols.

**Figure S8.** MethylFLYA performance on different input structures for three enzymes in the benchmark.

**Supplementary Methods**

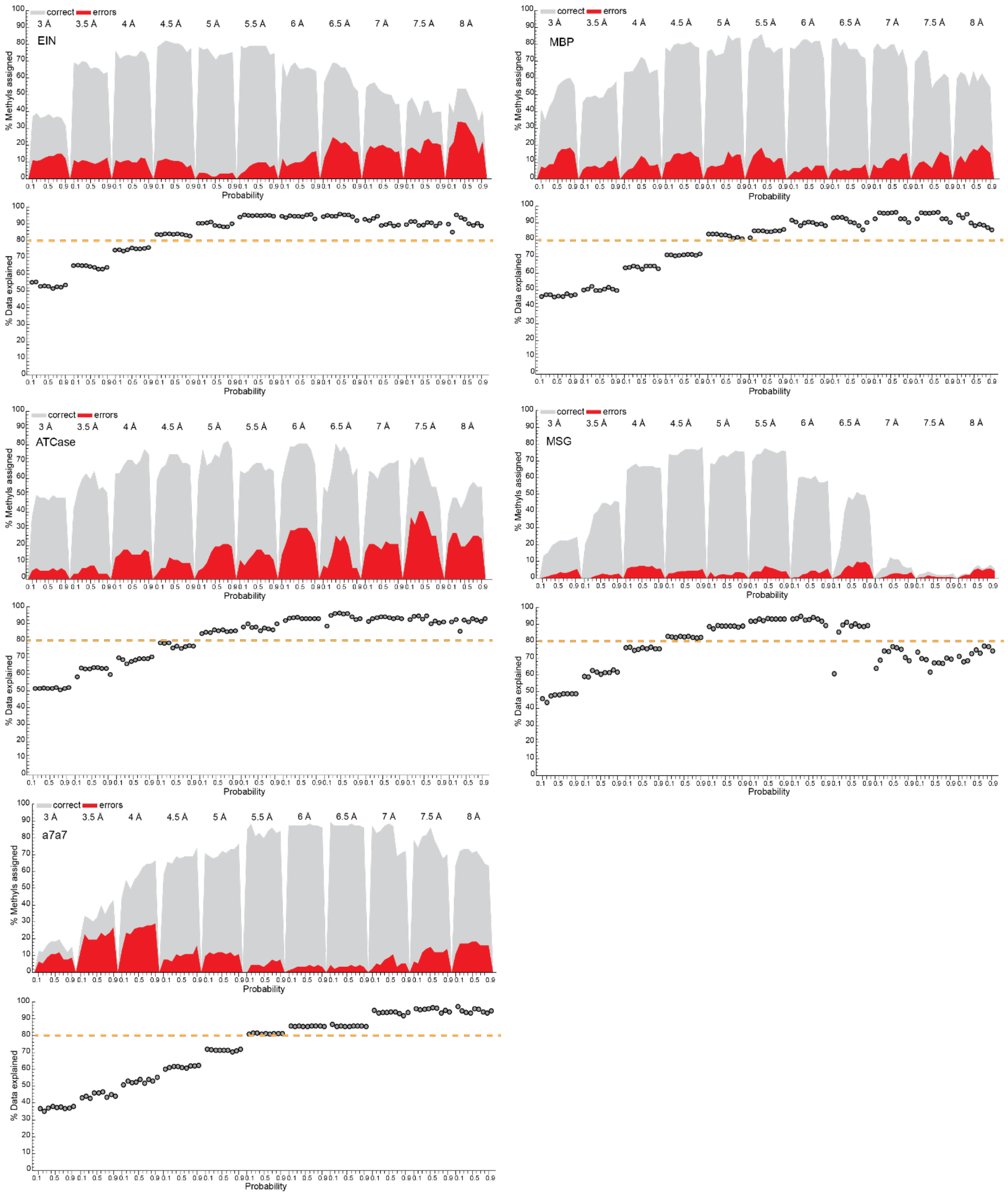

**Figure S1.** Parametrization of the distance cutoff $d_{cut}$ and the probability of expected methyl-methyl NOE contacts, $p_{NOE}$. For each protein, the upper panel shows the percentage of correct (*grey*) and erroneous (*red*) strong (i.e. confident) assignments for given $d_{cut}$ and $p_{NOE}$ values. Assignment percentages are relative to the number of reference assignments. For each protein, the lower panels show for each protein the percentage of explained experimental NOESY peaks. The distances at which ~80–85% of the data are explained generally led to the most reliable assignments. For the largest dataset (MSG) with >250 methyls, the algorithmic performance significantly deteriorates for $d_{cut}$ > 6.5 Å (equivalent to >11.5 Å C–C distance), resulting in an unreliable assignment. Note that such large distance cutoffs are unrealistic in practice and unlikely to be generally required.

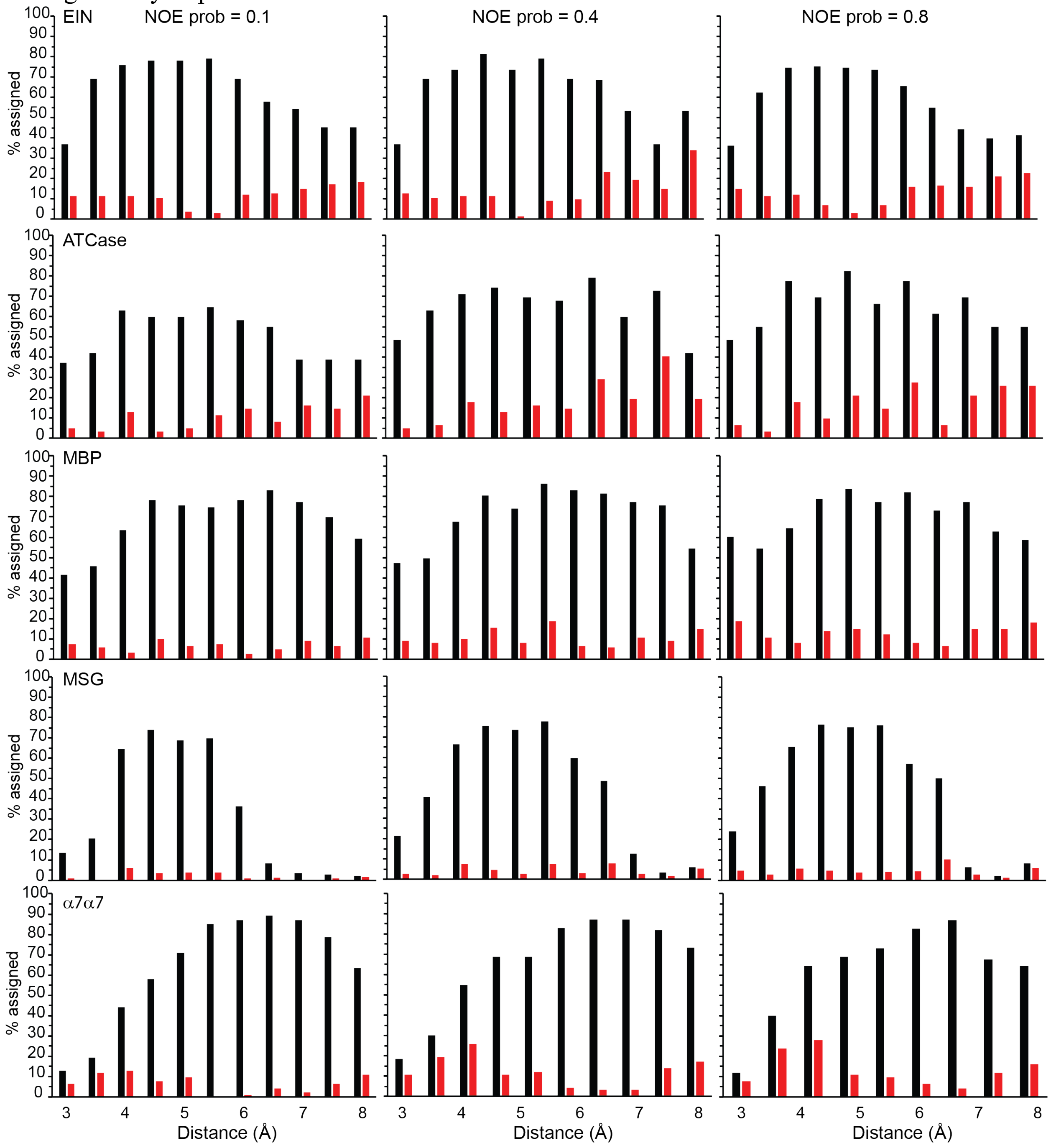

**Figure S2.** Accuracy of the methyl assignments obtained for the different values of NOE probabilities over a range of NOE distance cutoffs, $d_{cut}$. The percentage of accurately (*black*) and erroneously (*red*) assigned methyl groups is shown for the NOE probability values of 0.1, 0.4, and 0.8 in the first, second, and third column, respectively.

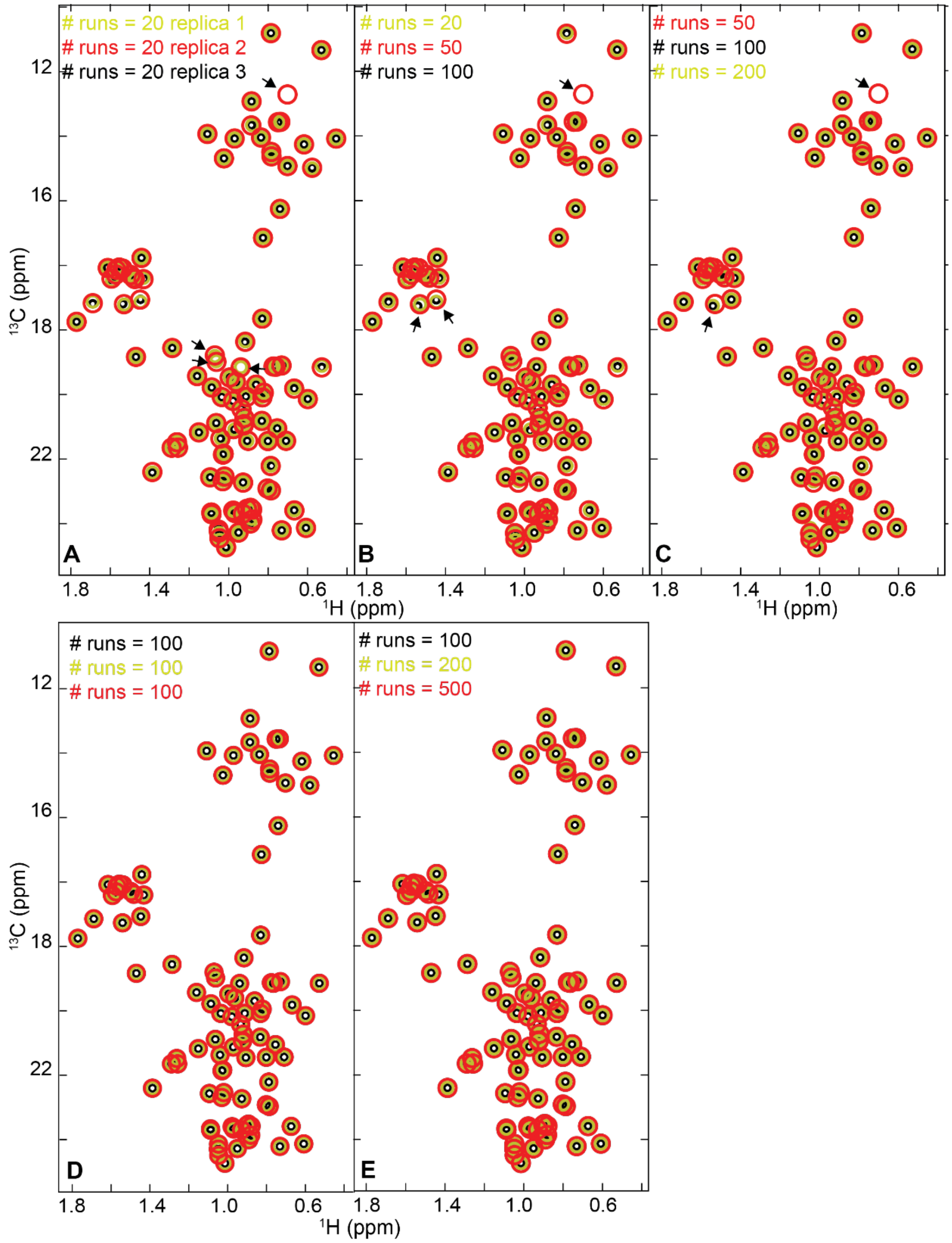

**Figure S3.** Determination of the optimal number of individual assignment runs for automatic methyl resonance assignment with MethylFLYA. Positions of the "strong" (i.e. confident) MethylFLYA-derived methyl assignments for EIN are shown in $^{1}$H-$^{13}$C correlation plots as circles with different colors and increasing diameters. **(A)** Running 20 parallel assignment calculations in three replicates, each from a different random starting point, shows some differences in the derived strong assignments between the replicates (arrows). **(B-C)** Increasing the number of calculations to 50 shows that the differences persist when compared to the higher number of parallel calculations (100 or 200). **(D)** Running 100 parallel MethylFLYA calculations is sufficient for the reproducibility of strong assignments. **(E)** A further increase in the number of parallel calculations (e.g. 200, 500) results in sets of strong assignments that are consistent with the set of 100 parallel calculations.

**Table S1.** Methyl resonance assignments by MethylFLYA, MAGMA, MAP-XSII, and FLAMEnGO2.0. The methyl groups with reference assignment are listed, with the $^{13}C$ and $^{1}H$ chemical shift values in ppm in columns '13C' and '1H'. Agreement or disagreement of the confident assignments by the four algorithms with the reference assignment is indicated by '=' (agreement) or '!' (disagreement) signs. The first sign is for $^{13}C$, the second for $^{1}H$. Missing, non-confident, or ambiguous assignments are left blank, except in the case of MethylFLYA, where also the weak (non-confident) assignments are indicated in parentheses. Filtered peak lists were used.

ATCase

| Methyl | 13C | 1H | FLYA | MAGMA | MAPXS | FLAMENGO |
|---|---|---|---|---|---|---|
| 7 LEU CD1 | 24.700 | 0.825 | == | | | |
| 7 LEU CD2 | 23.351 | 0.779 | == | | | |
| 9 VAL CG1 | 20.219 | 0.870 | == | | | |
| 9 VAL CG2 | 21.153 | 0.879 | == | | | |
| 12 ILE CD1 | 13.256 | 0.486 | (==) | | !! | == |
| 17 VAL CG1 | 20.691 | 0.894 | (==) | | | |
| 18 ILE CD1 | 13.418 | 0.734 | == | | == | |
| 21 ILE CD1 | 14.203 | 0.678 | (==) | | == | |
| 30 LEU CD1 | 25.344 | 0.851 | == | == | | |
| 30 LEU CD2 | 23.065 | 0.650 | == | == | | |
| 32 LEU CD1 | 23.190 | 0.364 | == | | | == |
| 32 LEU CD2 | 24.650 | 0.286 | == | | | == |
| 35 LEU CD1 | 24.537 | 0.727 | == | == | | |
| 35 LEU CD2 | 21.752 | 0.733 | == | == | | |
| 42 ILE CD1 | 14.481 | 0.676 | !! | == | == | |
| 44 ILE CD1 | 16.259 | 0.769 | !! | == | == | |
| 46 LEU CD2 | 22.309 | 0.823 | == | == | | !! |
| 48 LEU CD1 | 22.510 | 0.844 | (!!) | | | |
| 48 LEU CD2 | 27.064 | 0.917 | (!!) | | | |
| 58 LEU CD1 | 24.286 | 0.781 | (==) | | == | |
| 58 LEU CD2 | 26.475 | 0.732 | (==) | | | |
| 59 ILE CD1 | 14.045 | 0.754 | =! | | == | |
| 61 ILE CD1 | 14.380 | 0.679 | =! | | == | |
| 66 LEU CD1 | 24.699 | 0.617 | (==) | | !! | |
| 66 LEU CD2 | 21.671 | 0.481 | (==) | | | |
| 71 VAL CG1 | 21.088 | 0.790 | == | | | |
| 71 VAL CG2 | 23.701 | 0.902 | == | | | |

| Methyl | 13C | 1H | FLYA | MAGMA | MAPXS | FLAMENGO |
|---|---|---|---|---|---|---|
| 74 LEU CD1 | 25.859 | 0.789 | (==) | | !! | !! |
| 76 LEU CD1 | 25.036 | 0.788 | == | | == | |
| 76 LEU CD2 | 24.234 | 0.659 | == | | == | |
| 83 VAL CG1 | 21.482 | 0.783 | == | | | |
| 83 VAL CG2 | 21.844 | 0.779 | == | | | |
| 86 ILE CD1 | 11.600 | 0.602 | (==) | | | == |
| 91 VAL CG1 | 21.777 | 0.944 | == | | == | |
| 91 VAL CG2 | 21.184 | 0.954 | == | | == | |
| 92 VAL CG1 | 21.705 | 1.032 | == | | | |
| 92 VAL CG2 | 18.600 | 0.761 | == | | == | |
| 99 LEU CD1 | 25.443 | 0.739 | (==) | == | == | |
| 99 LEU CD2 | 24.463 | 0.751 | (==) | == | == | |
| 103 ILE CD1 | 13.711 | 0.794 | == | == | == | == |
| 106 VAL CG2 | 19.857 | 0.939 | == | | == | |
| 107 LEU CD1 | 25.746 | 0.448 | == | | | |
| 107 LEU CD2 | 21.905 | 0.533 | == | | | |
| 127 VAL CG1 | 21.790 | 0.696 | == | | == | !! |
| 127 VAL CG2 | 20.699 | 0.620 | == | | == | == |
| 134 ILE CD1 | 13.073 | 0.492 | == | == | == | == |
| 136 LEU CD1 | 25.551 | 0.682 | (==) | | == | |
| 136 LEU CD2 | 25.695 | 0.588 | (==) | | == | |
| 149 VAL CG1 | 21.889 | 1.161 | == | | | |
| 149 VAL CG2 | 20.361 | 0.954 | == | | == | |
| 150 VAL CG1 | 22.056 | 0.591 | == | | == | !! |
| 150 VAL CG2 | 22.442 | 0.733 | == | | == | == |
| 151 LEU CD1 | 23.053 | 0.512 | == | | | |
| 151 LEU CD2 | 24.279 | 0.458 | == | | | |

EIN

| Methyl | 13C | 1H | FLYA | MAGMA | MAPXS | FLAMENGO |
|---|---|---|---|---|---|---|
| 6 LEU CD1 | 23.060 | 0.755 | == | == | | |
| 6 LEU CD2 | 25.480 | 0.891 | == | == | | |
| 11 ILE CD1 | 14.030 | 0.834 | == | == | == | |
| 12 ALA CB | 23.660 | 1.258 | == | == | == | |
| 16 ALA CB | 20.550 | 1.287 | == | == | == | |
| 17 LEU CD1 | 25.850 | 0.884 | == | | !! | |
| 17 LEU CD2 | 23.840 | 1.025 | == | | !! | |
| 18 LEU CD1 | 25.640 | 0.978 | == | == | !! | |
| 18 LEU CD2 | 23.450 | 0.800 | == | == | | |
| 19 LEU CD1 | 25.500 | 0.963 | (==) | | !! | |
| 19 LEU CD2 | 23.980 | 0.890 | (==) | | !! | |
| 24 ILE CD1 | 14.070 | 0.970 | == | == | == | |
| 26 ILE CD1 | 11.330 | 0.531 | == | == | == | |
| 31 ILE CD1 | 14.940 | 0.697 | == | == | == | |
| 33 ALA CB | 18.340 | 1.573 | == | == | | |
| 36 VAL CG1 | 21.790 | 1.087 | == | | | |
| 36 VAL CG2 | 23.440 | 1.255 | == | | == | |
| 40 VAL CG1 | 21.440 | 1.161 | == | | == | |
| 40 VAL CG2 | 23.690 | 1.294 | == | | == | |
| 44 LEU CD1 | 21.810 | 0.669 | == | == | | |
| 44 LEU CD2 | 25.730 | 0.927 | == | == | | |
| 50 ALA CB | 18.380 | 1.482 | == | | | |
| 54 LEU CD1 | 26.170 | 1.046 | == | == | | |
| 54 LEU CD2 | 23.180 | 1.151 | (==) | == | | |
| 57 ILE CD1 | 13.640 | 0.742 | (=!) | | !! | |
| 61 ALA CB | 18.520 | 1.593 | (==) | | !! | |
| 71 ALA CB | 18.220 | 1.530 | (!!) | | == | |
| 72 ILE CD1 | 12.740 | 0.697 | == | | == | |
| 77 ILE CD1 | 13.390 | 0.888 | (==) | | | |
| 79 LEU CD1 | 23.460 | 0.903 | == | | | |
| 79 LEU CD2 | 25.500 | 0.916 | == | | | |
| 80 LEU CD1 | 26.580 | 1.046 | == | | | |
| 80 LEU CD2 | 24.410 | 1.011 | == | | | |

| Methyl | 13C | 1H | FLYA | MAGMA | MAPXS | FLAMENGO |
|---|---|---|---|---|---|---|
| 137 LEU CD1 | 22.080 | 1.036 | == | | == | |
| 137 LEU CD2 | 25.720 | 1.088 | == | | | |
| 138 LEU CD1 | 25.620 | 1.089 | == | == | == | |
| 138 LEU CD2 | 24.580 | 1.095 | == | == | == | |
| 141 ILE CD1 | 14.690 | 1.022 | == | == | == | |
| 142 LEU CD1 | 26.290 | 0.949 | == | == | == | |
| 142 LEU CD2 | 22.890 | 1.063 | == | == | | |
| 147 ILE CD1 | 12.910 | 0.884 | (==) | == | == | |
| 149 LEU CD2 | 23.210 | 0.850 | == | | !! | |
| 151 ALA CB | 19.030 | 1.427 | (==) | == | == | |
| 152 ILE CD1 | 13.680 | 0.883 | == | == | == | |
| 156 VAL CG1 | 22.100 | 0.832 | == | | !! | |
| 156 VAL CG2 | 17.160 | 0.826 | == | | !! | |
| 157 ILE CD1 | 14.070 | 0.457 | == | == | == | |
| 158 LEU CD1 | 25.840 | 0.976 | == | | == | |
| 159 VAL CG1 | 23.450 | 0.710 | == | | == | |
| 159 VAL CG2 | 21.190 | 0.761 | == | | == | |
| 160 ALA CB | 24.410 | 1.387 | == | == | == | |
| 161 ALA CB | 18.120 | 1.617 | == | | == | |
| 163 LEU CD1 | 25.570 | 0.670 | (==) | | | |
| 163 LEU CD2 | 24.230 | 0.782 | (==) | | | |
| 169 ALA CB | 19.220 | 1.459 | (==) | | !! | |
| 176 VAL CG1 | 22.150 | 0.600 | == | | !! | |
| 176 VAL CG2 | 22.060 | 0.914 | == | | !! | |
| 177 LEU CD1 | 26.030 | 0.889 | == | == | == | |
| 177 LEU CD2 | 21.700 | 0.860 | == | == | | |
| 180 ILE CD1 | 16.240 | 0.740 | == | == | == | |
| 183 ALA CB | 20.740 | 1.604 | (!!) | | == | |
| 194 ALA CB | 18.450 | 1.470 | (==) | | | |
| 197 LEU CD1 | 26.601 | 1.011 | == | | | |
| 199 LEU CD1 | 26.133 | 0.840 | (=!) | | | |
| 201 ALA CB | 22.220 | 1.031 | (!!) | | | |
| 202 ILE CD1 | 13.550 | 0.731 | == | == | == | |

| | | | | | |
|---|---|---|---|---|---|
| 85 LEU CD1 | 24.750 | 1.031 | == | | == |
| 85 LEU CD2 | 24.730 | 0.927 | == | | == |
| 89 ILE CD1 | 14.270 | 0.619 | == | == | == |
| 90 ILE CD1 | 13.940 | 1.107 | == | == | == |
| 91 ALA CB | 18.060 | 1.544 | == | == | |
| 92 LEU CD1 | 26.780 | 1.011 | == | == | |
| 92 LEU CD2 | 22.750 | 0.915 | == | == | |
| 93 ILE CD1 | 14.460 | 0.786 | == | == | == |
| 100 ALA CB | 19.750 | 1.771 | == | | == |
| 102 ALA CB | 18.270 | 1.496 | == | | == |
| 103 ALA CB | 20.840 | 1.470 | == | | |
| 104 ALA CB | 17.780 | 1.442 | == | | == |
| 107 VAL CG1 | 21.610 | 0.961 | == | | == |
| 107 VAL CG2 | 23.870 | 1.031 | == | | == |
| 108 ILE CD1 | 10.820 | 0.787 | == | == | == |
| 112 ALA CB | 18.440 | 1.429 | == | | == |
| 114 ALA CB | 18.000 | 1.540 | == | | |
| 115 LEU CD1 | 25.590 | 0.866 | == | == | == |
| 115 LEU CD2 | 23.090 | 0.976 | == | == | |
| 118 LEU CD2 | 22.450 | 0.945 | (!=) | | |
| 123 LEU CD1 | 26.150 | 1.062 | == | | |
| 127 ALA CB | 19.170 | 1.688 | == | | == |
| 130 VAL CG1 | 22.207 | 0.976 | == | | == |
| 130 VAL CG2 | 23.390 | 1.038 | == | | == |
| 133 ILE CD1 | 14.630 | 0.784 | == | == | == |

| | | | | | |
|---|---|---|---|---|---|
| 203 VAL CG1 | 19.700 | 0.729 | (==) | | |
| 208 VAL CG1 | 22.860 | 0.922 | == | | == |
| 208 VAL CG2 | 19.660 | 0.830 | == | | == |
| 212 VAL CG1 | 23.000 | 0.925 | == | | == |
| 212 VAL CG2 | 22.820 | 0.836 | == | | == |
| 218 LEU CD1 | 26.130 | 0.609 | == | == | == |
| 218 LEU CD2 | 24.900 | 0.806 | == | == | == |
| 219 ILE CD1 | 14.980 | 0.580 | == | | == |
| 220 LEU CD1 | 24.990 | 0.781 | == | == | == |
| 220 LEU CD2 | 26.220 | 0.731 | == | == | == |
| 222 ALA CB | 18.140 | 1.568 | == | == | == |
| 223 VAL CG1 | 20.380 | 0.918 | (==) | | |
| 223 VAL CG2 | 21.200 | 0.936 | (==) | | == |
| 227 VAL CG1 | 21.070 | 0.718 | == | | == |
| 227 VAL CG2 | 21.940 | 0.818 | == | | == |
| 229 VAL CG1 | 21.470 | 0.996 | == | | == |
| 229 VAL CG2 | 21.080 | 0.786 | == | | == |
| 235 VAL CG1 | 21.160 | 0.527 | == | | == |
| 236 ILE CD1 | 13.530 | 0.754 | == | | == |
| 241 ALA CB | 17.870 | 1.594 | (==) | | |
| 242 VAL CG1 | 21.020 | 0.981 | (=!) | | !! |
| 242 VAL CG2 | 22.040 | 1.167 | (!) | | |
| 246 VAL CG1 | 21.100 | 1.041 | (==) | | |
| 246 VAL CG2 | 20.840 | 1.075 | (==) | | |
| 247 ALA CB | 19.240 | 1.534 | (==) | | |

MBP

| Methyl | 13C | 1H | FLYA | MAGMA | MAPXS | FLAMENGO |
|---|---|---|---|---|---|---|
| 2 ILE CD1 | 13.054 | 0.402 | (=!) | == | !! | |
| 7 LEU CD1 | 25.203 | 0.267 | == | == | !! | == |
| 7 LEU CD2 | 24.064 | 0.615 | == | == | | == |
| 8 VAL CG1 | 20.962 | 0.876 | (!) | | | |
| 8 VAL CG2 | 21.211 | 0.923 | (=!) | | | |
| 9 ILE CD1 | 14.523 | 0.346 | == | == | !! | == |
| 11 ILE CD1 | 13.105 | 0.107 | == | == | !! | |
| 20 LEU CD1 | 24.725 | 0.876 | == | | | |
| 20 LEU CD2 | 26.879 | 0.638 | == | | !! | |
| 23 VAL CG1 | 22.314 | 0.823 | == | | | |
| 23 VAL CG2 | 23.403 | 1.252 | == | | | |
| 33 ILE CD1 | 10.138 | 0.574 | == | == | | == |
| 35 VAL CG1 | 20.685 | 0.331 | == | | | |
| 35 VAL CG2 | 22.360 | 0.760 | == | | | |
| 37 VAL CG1 | 22.840 | 0.951 | == | | | |
| 37 VAL CG2 | 20.673 | 0.766 | == | | | |
| 43 LEU CD1 | 26.677 | 1.101 | == | == | | |
| 43 LEU CD2 | 26.431 | 1.133 | == | == | !! | |
| 50 VAL CG1 | 21.260 | 0.967 | (!!) | | | |
| 50 VAL CG2 | 19.894 | 0.906 | (!!) | | | |
| 59 ILE CD1 | 13.790 | 0.735 | == | == | | |
| 60 ILE CD1 | 14.568 | 0.452 | == | == | == | |
| 75 LEU CD1 | 25.725 | 0.638 | (=!) | == | | |
| 75 LEU CD2 | 20.664 | 0.538 | (!!) | == | | |
| 76 LEU CD1 | 27.430 | 0.897 | == | == | | |
| 76 LEU CD2 | 21.891 | 0.538 | == | == | == | |
| 79 ILE CD1 | 12.313 | 0.161 | == | == | == | |
| 89 LEU CD1 | 26.152 | 0.526 | == | == | | |
| 89 LEU CD2 | 25.298 | 0.484 | == | == | | |
| 97 VAL CG1 | 22.085 | 0.578 | == | | | == |
| 97 VAL CG2 | 19.363 | 1.347 | == | | | == |
| 103 LEU CD1 | 24.748 | 0.490 | == | == | | |
| 103 LEU CD2 | 24.100 | 0.972 | == | == | | |
| 104 ILE CD1 | 14.779 | 0.845 | == | == | | |
| 108 ILE CD1 | 9.981 | 0.637 | == | == | == | |
| 110 VAL CG1 | 20.954 | 0.767 | == | | | |
| 110 VAL CG2 | 20.424 | 0.476 | == | | | |
| 115 LEU CD1 | 22.828 | 0.684 | (==) | | | |
| 115 LEU CD2 | 24.597 | 0.587 | (!) | | | |
| 116 ILE CD1 | 13.706 | 0.459 | (!!) | | | |
| 121 LEU CD1 | 24.729 | 0.665 | == | | | |
| 121 LEU CD2 | 22.791 | 0.779 | == | | | |
| 122 LEU CD1 | 25.548 | 0.930 | (!!) | | | |
| 122 LEU CD2 | 26.848 | 0.937 | (!!) | | | |
| 132 ILE CD1 | 12.360 | 0.560 | == | | | |
| 135 LEU CD1 | 24.040 | 0.888 | (!!) | | | |
| 135 LEU CD2 | 24.317 | 0.882 | (!=) | | | |
| 139 LEU CD1 | 25.914 | 0.787 | =! | | | |
| 139 LEU CD2 | 22.927 | 0.852 | !! | | | |
| 147 LEU CD1 | 22.500 | 0.683 | (==) | | | |

| Methyl | 13C | 1H | FLYA | MAGMA | MAPXS | FLAMENGO |
|---|---|---|---|---|---|---|
| 183 VAL CG1 | 22.416 | 0.882 | == | | | == |
| 183 VAL CG2 | 22.420 | 0.841 | == | | == | == |
| 192 LEU CD1 | 24.249 | 1.085 | == | == | == | == |
| 192 LEU CD2 | 25.330 | 0.845 | == | == | !! | == |
| 195 LEU CD1 | 23.706 | 0.893 | == | == | | |
| 195 LEU CD2 | 25.191 | 0.845 | == | == | | |
| 196 VAL CG1 | 21.649 | 0.969 | == | | | |
| 196 VAL CG2 | 23.319 | 0.856 | == | | | |
| 198 LEU CD1 | 25.598 | 1.018 | (!!) | | | |
| 198 LEU CD2 | 23.461 | 0.813 | (!!) | | | |
| 199 ILE CD1 | 14.222 | 0.743 | == | == | | == |
| 212 ILE CD1 | 13.110 | 0.867 | (==) | | | == |
| 226 ILE CD1 | 11.775 | 0.060 | == | | | == |
| 240 VAL CG1 | 20.890 | 0.585 | (=!) | | | |
| 240 VAL CG2 | 21.830 | 0.788 | (!!) | | | == |
| 244 VAL CG1 | 22.083 | 1.072 | (!!) | | | |
| 244 VAL CG2 | 21.606 | 0.636 | (=!) | | | |
| 246 VAL CG1 | 20.441 | 0.969 | == | | | |
| 246 VAL CG2 | 21.596 | 0.983 | == | | !! | |
| 247 LEU CD1 | 25.833 | 0.663 | (!=) | | | == |
| 247 LEU CD2 | 23.824 | 0.645 | (!!) | | | |
| 259 VAL CG1 | 21.509 | 0.925 | == | | | == |
| 259 VAL CG2 | 22.697 | 0.980 | == | | | == |
| 261 VAL CG1 | 20.308 | 0.679 | == | | | == |
| 261 VAL CG2 | 20.936 | 0.565 | == | | | |
| 262 LEU CD1 | 22.681 | 0.806 | == | == | | |
| 262 LEU CD2 | 25.314 | 0.885 | == | == | | |
| 266 ILE CD1 | 13.882 | 0.915 | == | == | | |
| 275 LEU CD1 | 25.202 | 0.933 | == | == | | == |
| 275 LEU CD2 | 22.493 | 0.947 | == | == | | == |
| 280 LEU CD1 | 25.779 | 0.672 | == | == | | == |
| 280 LEU CD2 | 23.405 | 0.710 | == | == | | |
| 284 LEU CD1 | 23.945 | 0.843 | == | | | |
| 284 LEU CD2 | 27.398 | 1.039 | == | | | |
| 285 LEU CD1 | 25.695 | 1.052 | == | == | | |
| 285 LEU CD2 | 24.003 | 1.119 | == | == | | |
| 290 LEU CD1 | 27.886 | 1.074 | == | == | | |
| 290 LEU CD2 | 21.864 | 0.792 | == | == | | |
| 293 VAL CG1 | 21.422 | 1.004 | == | | | |
| 293 VAL CG2 | 22.246 | 0.901 | == | | | |
| 302 VAL CG1 | 22.153 | 1.059 | == | | | |
| 302 VAL CG2 | 19.070 | 1.262 | == | | | |
| 304 LEU CD1 | 24.263 | 0.848 | (==) | == | | |
| 304 LEU CD2 | 26.586 | 0.702 | (==) | == | | |
| 311 LEU CD1 | 26.326 | 0.870 | == | | | == |
| 311 LEU CD2 | 22.652 | 0.797 | == | | | == |
| 317 ILE CD1 | 12.483 | 1.118 | (==) | == | | == |
| 329 ILE CD1 | 12.362 | 0.821 | == | == | == | == |
| 333 ILE CD1 | 12.253 | -0.138 | (==) | | == | == |
| 343 VAL CG1 | 21.297 | 0.863 | == | | == | == |

| | | | | | | |
|---|---|---|---|---|---|---|
| 147 LEU CD2 | 27.555 | 0.936 | (==) | | | |
| 151 LEU CD1 | 26.232 | 0.744 | == | | | |
| 151 LEU CD2 | 23.579 | 0.135 | == | | !! | |
| 160 LEU CD1 | 19.602 | -1.003 | (==) | == | == | |
| 160 LEU CD2 | 25.084 | -0.302 | (==) | == | !! | |
| 161 ILE CD1 | 13.081 | 0.760 | == | == | | == |
| 178 ILE CD1 | 13.905 | 0.853 | (==) | | == | == |
| 181 VAL CG1 | 21.539 | 0.823 | == | | == | == |
| 181 VAL CG2 | 22.538 | 0.711 | == | | | |
| 343 VAL CG2 | 23.236 | 0.168 | == | | == | == |
| 347 VAL CG1 | 20.468 | 0.918 | == | | | |
| 347 VAL CG2 | 22.282 | 1.043 | == | | | |
| 348 ILE CD1 | 12.965 | 0.869 | (==) | == | | == |
| 357 VAL CG1 | 21.315 | 0.991 | == | | | |
| 357 VAL CG2 | 23.401 | 1.045 | == | | !! | |
| 361 LEU CD1 | 25.264 | 0.892 | == | == | | == |
| 361 LEU CD2 | 19.605 | 0.722 | == | == | == | == |
| 368 ILE CD1 | 15.285 | 0.693 | == | == | == | == |

MSG

| Methyl | 13C | 1H | FLYA | MAGMA | MAPXS | FLAMENGO |
|---|---|---|---|---|---|---|
| 5 ILE CD1 | 12.833 | 0.820 | (==) | | | |
| 10 LEU CD1 | 24.486 | 0.915 | == | | | |
| 10 LEU CD2 | 22.364 | 0.868 | == | | | |
| 12 ILE CD1 | 15.537 | 0.008 | == | | | |
| 20 VAL CG1 | 21.566 | 0.463 | == | | | |
| 20 VAL CG2 | 23.955 | 0.739 | == | | | |
| 24 VAL CG1 | 20.604 | 0.541 | == | | | |
| 24 VAL CG2 | 22.556 | 0.137 | == | | | |
| 25 LEU CD1 | 24.621 | 0.295 | == | == | | |
| 25 LEU CD2 | 20.071 | -0.285 | == | == | | |
| 30 LEU CD1 | 25.838 | 0.755 | == | == | | |
| 30 LEU CD2 | 22.144 | 0.817 | == | == | | |
| 42 ILE CD1 | 14.704 | 0.680 | == | == | | |
| 43 VAL CG1 | 20.943 | 0.678 | == | | | |
| 43 VAL CG2 | 24.763 | 1.051 | == | | | |
| 46 LEU CD1 | 26.301 | 0.806 | (==) | | | |
| 46 LEU CD2 | 22.903 | 0.827 | (==) | | | |
| 53 LEU CD1 | 25.988 | 0.305 | (==) | | | |
| 54 LEU CD1 | 24.942 | 1.101 | == | | | |
| 54 LEU CD2 | 19.976 | 0.673 | == | | | |
| 60 ILE CD1 | 12.120 | 0.769 | == | == | | |
| 64 LEU CD1 | 25.428 | 0.904 | == | == | | |
| 64 LEU CD2 | 24.554 | 0.751 | == | == | | |
| 75 VAL CG2 | 20.637 | 0.815 | (==) | | | |
| 85 LEU CD1 | 24.801 | -0.085 | == | == | | |
| 85 LEU CD2 | 20.081 | -0.210 | == | == | | |
| 88 LEU CD1 | 23.547 | 0.823 | == | == | | |
| 88 LEU CD2 | 25.333 | 0.674 | == | == | | |
| 91 LEU CD1 | 25.353 | 0.901 | (!!) | | | |
| 91 LEU CD2 | 24.612 | 0.903 | (!!) | | | |
| 98 VAL CG1 | 20.318 | 0.852 | (==) | | | |
| 100 VAL CG1 | 21.311 | 0.958 | == | | | |
| 100 VAL CG2 | 22.218 | 0.857 | == | | | |
| 105 ILE CD1 | 10.255 | 0.810 | (!=) | !! | | |
| 109 ILE CD1 | 14.490 | 0.724 | (==) | !! | | |
| 117 LEU CD1 | 25.348 | 0.578 | == | | | |
| 117 LEU CD2 | 26.534 | 0.593 | == | | | |
| 118 VAL CG1 | 19.382 | 0.230 | | | | |
| 118 VAL CG2 | 17.697 | -0.834 | (!) | | !! | |
| 119 VAL CG1 | 21.619 | 0.737 | == | | | |
| 119 VAL CG2 | 18.891 | 0.687 | == | | | |
| 128 LEU CD1 | 24.941 | 0.572 | (==) | | | |
| 128 LEU CD2 | 24.322 | 0.607 | (==) | | | |
| 138 LEU CD1 | 23.192 | 0.768 | (==) | | | |
| 138 LEU CD2 | 26.304 | 0.748 | (==) | | | |
| 142 LEU CD1 | 27.936 | 0.456 | == | == | | |
| 142 LEU CD2 | 21.862 | -0.448 | == | == | | |
| 147 ILE CD1 | 13.610 | 0.262 | == | == | | |
| 148 ILE CD1 | 11.341 | 0.390 | == | == | | |
| 166 VAL CG1 | 21.207 | 0.848 | == | | | |
| 166 VAL CG2 | 22.805 | 0.924 | == | | | |
| 167 ILE CD1 | 14.215 | 0.831 | (!!) | | | |
| 170 VAL CG1 | 22.279 | 1.178 | == | | == | |
| 170 VAL CG2 | 25.567 | 1.376 | == | | | |
| 178 LEU CD1 | 26.784 | 0.611 | == | | == | |
| 178 LEU CD2 | 26.758 | 0.726 | == | | == | |
| 180 LEU CD1 | 24.390 | -0.082 | == | == | | |
| 180 LEU CD2 | 22.737 | 0.359 | == | == | | |
| 188 VAL CG1 | 21.750 | 0.407 | == | | | |
| 188 VAL CG2 | 23.700 | 0.743 | == | | | |
| 189 VAL CG2 | 18.759 | 0.541 | == | | | |
| 193 VAL CG1 | 19.420 | 0.703 | == | | | |
| 193 VAL CG2 | 19.366 | 0.205 | == | | | |
| 194 VAL CG2 | 20.321 | 0.773 | (=!) | | | |
| 198 LEU CD1 | 22.639 | 0.663 | == | == | | |
| 198 LEU CD2 | 26.102 | 0.813 | == | == | | |

| Methyl | 13C | 1H | FLYA | MAGMA | MAPXS | FLAMENGO |
|---|---|---|---|---|---|---|
| 348 VAL CG1 | 20.967 | 0.163 | == | | | |
| 348 VAL CG2 | 21.448 | 0.402 | == | | | |
| 349 ILE CD1 | 14.919 | 0.778 | == | | | |
| 361 ILE CD1 | 12.815 | 0.999 | == | | !! | |
| 362 LEU CD1 | 24.066 | 1.105 | == | | | |
| 362 LEU CD2 | 27.381 | 0.769 | == | | | |
| 365 VAL CG1 | 20.364 | -0.085 | == | | | |
| 365 VAL CG2 | 21.867 | 0.566 | == | | | |
| 370 ILE CD1 | 13.820 | 0.871 | == | == | | |
| 372 LEU CD1 | 23.064 | 0.863 | == | == | | |
| 372 LEU CD2 | 25.825 | 1.033 | == | == | !! | |
| 375 LEU CD1 | 25.628 | 0.885 | == | == | | |
| 375 LEU CD2 | 22.069 | 0.721 | == | == | | |
| 386 VAL CG1 | 19.406 | 0.670 | == | | | |
| 386 VAL CG2 | 21.071 | 0.726 | == | | | |
| 388 ILE CD1 | 12.747 | 0.290 | == | == | | |
| 389 VAL CG1 | 22.383 | 0.720 | (==) | | | |
| 389 VAL CG2 | 20.605 | 0.437 | (==) | | | |
| 399 VAL CG1 | 21.973 | 0.912 | == | | | |
| 399 VAL CG2 | 20.715 | 0.989 | == | | | |
| 409 ILE CD1 | 13.073 | 0.548 | == | == | | |
| 413 LEU CD1 | 26.081 | 0.718 | == | == | | |
| 413 LEU CD2 | 22.095 | 0.807 | == | == | | |
| 420 LEU CD1 | 24.802 | 0.469 | == | == | | |
| 420 LEU CD2 | 22.504 | 0.373 | == | == | | |
| 424 ILE CD1 | 14.922 | 0.745 | == | == | == | |
| 433 LEU CD1 | 27.191 | 0.971 | == | | | |
| 433 LEU CD2 | 25.452 | 0.888 | == | | | |
| 435 LEU CD1 | 27.524 | 1.194 | == | == | | |
| 435 LEU CD2 | 21.476 | 0.801 | == | == | == | |
| 439 ILE CD1 | 13.782 | 0.790 | == | | !! | |
| 446 VAL CG1 | 21.292 | 0.980 | == | | | |
| 446 VAL CG2 | 22.836 | 1.134 | == | | | |
| 449 ILE CD1 | 14.145 | 0.642 | == | | !! | |
| 464 VAL CG1 | 20.503 | 0.474 | == | | | |
| 464 VAL CG2 | 17.943 | 0.670 | == | | | |
| 471 LEU CD2 | 22.194 | 1.075 | (==) | == | | |
| 490 VAL CG1 | 19.622 | 0.904 | == | | | |
| 490 VAL CG2 | 20.949 | 0.854 | == | | | |
| 491 LEU CD1 | 25.082 | 0.561 | == | == | | |
| 491 LEU CD2 | 20.686 | 0.282 | == | == | | |
| 494 LEU CD1 | 25.945 | 0.627 | == | == | | |
| 494 LEU CD2 | 22.337 | 0.670 | == | == | | |
| 498 LEU CD1 | 25.082 | 0.761 | == | == | == | |
| 498 LEU CD2 | 23.324 | 0.619 | == | == | == | |
| 504 ILE CD1 | 13.295 | 0.722 | == | | | |
| 514 LEU CD1 | 25.980 | 1.066 | == | | | |
| 514 LEU CD2 | 22.371 | 0.937 | == | | | |
| 526 LEU CD1 | 23.498 | 0.115 | (==) | | | |
| 526 LEU CD2 | 22.406 | 0.100 | (==) | | == | |
| 535 VAL CG2 | 17.974 | 0.916 | == | | | |
| 543 LEU CD1 | 25.180 | 0.572 | == | == | | |
| 543 LEU CD2 | 21.491 | 0.250 | == | == | | |
| 546 LEU CD1 | 25.237 | 0.781 | (==) | | | |
| 546 LEU CD2 | 20.492 | 0.261 | (==) | | | |
| 553 VAL CG1 | 22.981 | 1.087 | (==) | | | |
| 553 VAL CG2 | 20.128 | 1.115 | (==) | | | |
| 556 VAL CG1 | 21.888 | 0.894 | (==) | | | |
| 556 VAL CG2 | 20.935 | 0.865 | (!=) | | | |
| 560 ILE CD1 | 13.815 | 0.603 | == | == | | |
| 572 LEU CD1 | 26.490 | 0.721 | == | == | | |
| 572 LEU CD2 | 23.384 | 1.055 | == | == | | |
| 573 LEU CD1 | 24.348 | 0.989 | == | | | |
| 573 LEU CD2 | 25.639 | 0.950 | == | | | |
| 576 LEU CD1 | 24.877 | 0.826 | == | == | | |
| 576 LEU CD2 | 24.663 | 1.214 | == | == | | |

| | | | | | |
|---|---|---|---|---|---|
| 200 ILE CD1 | 16.119 | 0.746 | == | == | == |
| 202 LEU CD1 | 24.801 | 0.768 | == | == | |
| 202 LEU CD2 | 22.680 | 0.501 | == | == | |
| 210 LEU CD1 | 28.067 | 1.090 | == | | == |
| 210 LEU CD2 | 22.536 | 0.904 | == | | == |
| 217 VAL CG1 | 21.989 | 0.800 | == | | |
| 217 VAL CG2 | 21.929 | 0.880 | == | | |
| 229 ILE CD1 | 10.616 | 0.562 | == | == | == |
| 230 LEU CD1 | 21.754 | 0.680 | == | | |
| 230 LEU CD2 | 24.072 | -0.499 | (==) | | |
| 231 LEU CD1 | 25.413 | 0.876 | == | | |
| 231 LEU CD2 | 24.834 | 1.069 | == | | |
| 236 LEU CD1 | 25.156 | 0.804 | (==) | == | |
| 236 LEU CD2 | 22.500 | 0.699 | (==) | == | |
| 238 ILE CD1 | 13.747 | 0.503 | == | == | == |
| 240 LEU CD1 | 24.913 | 0.632 | == | == | == |
| 240 LEU CD2 | 24.407 | 0.715 | == | == | |
| 242 ILE CD1 | 12.365 | 0.664 | == | | == |
| 248 ILE CD1 | 12.276 | 0.756 | (!!) | | |
| 256 ILE CD1 | 11.691 | 0.901 | == | | == |
| 259 VAL CG1 | 19.992 | 0.651 | == | | |
| 259 VAL CG2 | 21.191 | 0.648 | == | | == |
| 260 ILE CD1 | 11.138 | 0.651 | (==) | | !! |
| 261 VAL CG1 | 22.279 | 0.898 | == | | |
| 261 VAL CG2 | 21.020 | 0.924 | == | | |
| 265 ILE CD1 | 9.490 | 0.378 | (==) | == | |
| 268 ILE CD1 | 13.602 | 0.655 | (==) | | !! |
| 269 LEU CD1 | 26.885 | 0.876 | == | | |
| 269 LEU CD2 | 22.419 | 0.655 | == | | |
| 275 VAL CG1 | 23.138 | 0.833 | (==) | | |
| 275 VAL CG2 | 21.430 | 0.785 | (==) | | |
| 278 VAL CG1 | 21.032 | 0.423 | (==) | | |
| 278 VAL CG2 | 18.142 | 0.642 | (==) | | |
| 284 ILE CD1 | 13.198 | 0.745 | (==) | | |
| 286 LEU CD1 | 25.696 | 0.881 | (==) | | |
| 286 LEU CD2 | 26.157 | 0.768 | (==) | | |
| 291 LEU CD1 | 25.289 | 0.601 | == | == | |
| 291 LEU CD2 | 26.260 | 0.771 | == | == | |
| 293 LEU CD1 | 25.478 | 0.930 | (=!) | | |
| 293 LEU CD2 | 24.044 | 0.778 | (!=) | | |
| 298 LEU CD1 | 23.575 | 1.052 | (=!) | | |
| 298 LEU CD2 | 25.239 | 0.821 | (==) | | |
| 313 LEU CD1 | 25.130 | 0.938 | (=!) | | |
| 313 LEU CD2 | 24.015 | 0.807 | (==) | | |
| 327 ILE CD1 | 13.073 | 0.572 | == | | |
| 329 LEU CD1 | 25.272 | 0.204 | (==) | | |
| 329 LEU CD2 | 22.249 | 0.625 | == | | |
| 334 LEU CD2 | 27.669 | 0.780 | (==) | == | |
| 335 LEU CD1 | 25.213 | 0.818 | (==) | | |
| 335 LEU CD2 | 22.086 | 0.633 | (!!) | | |
| 337 ILE CD1 | 14.606 | 0.875 | == | == | |
| 343 LEU CD1 | 22.988 | 0.988 | == | | |
| 343 LEU CD2 | 25.785 | 0.745 | == | | |
| 577 LEU CD1 | 26.739 | 0.781 | (!!) | | |
| 577 LEU CD2 | 22.528 | 0.738 | (!) | | |
| 579 ILE CD1 | 14.656 | 0.757 | == | == | == |
| 581 VAL CG1 | 20.609 | 0.530 | == | | |
| 581 VAL CG2 | 20.334 | 0.527 | == | | |
| 592 ILE CD1 | 13.646 | 0.825 | == | | == |
| 596 LEU CD1 | 26.922 | 1.050 | == | | == |
| 596 LEU CD2 | 24.904 | 1.053 | == | | |
| 600 VAL CG1 | 22.462 | 0.752 | == | | == |
| 600 VAL CG2 | 25.068 | 0.974 | == | | |
| 603 ILE CD1 | 14.352 | 0.502 | (==) | | == |
| 604 LEU CD1 | 27.761 | 0.912 | == | == | |
| 604 LEU CD2 | 23.071 | 0.754 | == | == | |
| 607 VAL CG1 | 23.104 | 1.033 | == | | == |
| 607 VAL CG2 | 26.709 | 1.120 | == | | == |
| 608 VAL CG1 | 20.748 | 1.329 | (==) | | |
| 608 VAL CG2 | 21.975 | 1.155 | (==) | | |
| 611 VAL CG1 | 21.774 | 0.864 | == | | |
| 611 VAL CG2 | 25.146 | 1.349 | == | | == |
| 623 ILE CD1 | 14.613 | 0.784 | (==) | | |
| 626 VAL CG1 | 20.784 | 0.805 | == | | |
| 626 VAL CG2 | 21.600 | 0.882 | == | | |
| 628 LEU CD1 | 25.544 | 0.894 | == | | |
| 628 LEU CD2 | 23.460 | 0.678 | == | | |
| 635 LEU CD1 | 27.517 | 0.867 | == | | |
| 635 LEU CD2 | 25.306 | 1.020 | == | | |
| 642 ILE CD1 | 15.941 | 0.647 | == | | == |
| 646 LEU CD1 | 23.659 | 0.863 | == | | |
| 646 LEU CD2 | 25.235 | 0.829 | == | | |
| 650 ILE CD1 | 8.057 | 0.581 | == | | == |
| 651 LEU CD1 | 28.466 | 1.193 | == | | == |
| 651 LEU CD2 | 25.318 | 1.057 | == | | == |
| 656 VAL CG1 | 22.427 | 0.763 | == | | == |
| 656 VAL CG2 | 24.262 | 1.119 | == | | == |
| 660 LEU CD1 | 25.900 | 0.717 | == | == | |
| 660 LEU CD2 | 23.362 | 0.479 | == | == | |
| 666 VAL CG1 | 19.705 | 0.899 | == | | |
| 666 VAL CG2 | 21.958 | 1.127 | (==) | | |
| 667 VAL CG1 | 22.628 | 1.058 | == | | |
| 667 VAL CG2 | 23.404 | 1.131 | == | | |
| 696 LEU CD1 | 26.693 | 0.738 | == | | |
| 696 LEU CD2 | 24.144 | 0.903 | == | | |
| 697 ILE CD1 | 14.757 | 0.574 | (==) | | == |
| 699 LEU CD1 | 25.034 | 0.907 | == | | |
| 699 LEU CD2 | 22.156 | 0.881 | == | | |
| 711 LEU CD1 | 25.811 | 0.806 | == | | |
| 711 LEU CD2 | 22.836 | 1.022 | == | | |
| 712 LEU CD1 | 24.727 | 0.182 | == | | |
| 712 LEU CD2 | 22.944 | 0.057 | == | | !! |
| 717 LEU CD1 | 25.176 | 0.980 | (==) | | |
| 717 LEU CD2 | 23.695 | 0.956 | (==) | | |

a7a7

| Methyl | 13C | 1H | FLYA | MAGMA | MAPXS | FLAMENGO |
|---|---|---|---|---|---|---|
| 21 LEU CD1 | 25.996 | 0.607 | (=!) | == | == | == |
| 24 VAL CG1 | 24.215 | 0.930 | == | | | == |
| 24 VAL CG2 | 21.800 | 0.873 | == | | == | == |
| 31 VAL CG1 | 21.101 | 0.975 | == | | == | == |
| 38 LEU CD1 | 26.082 | 0.639 | == | == | | == |
| 38 LEU CD2 | 28.507 | 0.701 | == | == | == | == |
| 46 VAL CG1 | 19.945 | 0.821 | == | | == | == |
| 46 VAL CG2 | 22.010 | 0.938 | == | | == | == |
| 47 LEU CD1 | 25.259 | 0.714 | == | == | == | == |
| 47 LEU CD2 | 28.341 | 0.572 | == | == | == | !! |
| 48 LEU CD1 | 26.717 | 0.608 | == | == | == | == |
| 48 LEU CD2 | 23.970 | 0.748 | == | == | == | !! |
| 49 ILE CD1 | 14.168 | 0.594 | == | == | == | == |
| 54 VAL CG1 | 20.875 | 0.923 | == | | == | == |
| 54 VAL CG2 | 21.440 | 0.918 | == | | == | == |
| 58 LEU CD1 | 21.343 | 0.440 | (!!) | == | | == |
| 58 LEU CD2 | 26.121 | 0.784 | (=!) | == | == | == |
| 59 ILE CD1 | 13.464 | 0.791 | (==) | == | == | == |
| 64 ILE CD1 | 12.983 | 0.799 | == | == | == | == |
| 67 ILE CD1 | 13.905 | 0.542 | == | == | == | == |
| 69 LEU CD1 | 24.774 | 0.755 | == | == | == | == |
| 69 LEU CD2 | 22.438 | 0.425 | == | == | == | == |
| 113 VAL CG1 | 20.996 | 0.301 | == | | == | == |
| 113 VAL CG2 | 21.787 | -0.033 | == | | == | == |
| 116 VAL CG1 | 22.011 | 0.950 | == | | == | !! |
| 116 VAL CG2 | 21.613 | 0.940 | == | | == | !! |
| 134 VAL CG1 | 22.670 | 0.742 | == | | == | == |
| 134 VAL CG2 | 23.838 | 0.872 | == | | == | == |
| 136 LEU CD1 | 27.678 | 0.667 | == | == | == | !! |
| 136 LEU CD2 | 26.611 | 0.410 | == | == | == | !! |
| 137 ILE CD1 | 14.071 | 0.374 | == | == | == | == |
| 141 ILE CD1 | 12.458 | 0.616 | == | == | == | == |
| 144 ILE CD1 | 11.757 | 0.762 | == | == | | == |
| 148 LEU CD1 | 25.245 | 0.731 | == | == | == | == |
| 148 LEU CD2 | 26.003 | 0.454 | == | == | == | == |
| 157 ILE CD1 | 13.663 | 0.638 | (==) | == | == | == |
| 165 ILE CD1 | 13.629 | 0.425 | == | == | == | == |
| 172 VAL CG1 | 23.655 | 0.696 | == | | == | == |
| 172 VAL CG2 | 22.860 | 0.799 | == | | == | == |
| 173 VAL CG1 | 21.461 | 0.946 | == | | == | == |
| 173 VAL CG2 | 23.532 | 0.982 | == | | == | == |
| 176 LEU CD1 | 25.759 | 0.638 | (==) | == | == | == |
| 176 LEU CD2 | 22.386 | 0.901 | (==) | == | == | == |
| 184 LEU CD1 | 25.965 | 0.733 | == | == | == | == |

| | | | | | | | | |
|---|---|---|---|---|---|---|---|---|
| 70 | ILE | CD1 | 11.698 | 0.660 | == | == | == | == |
| 74 | VAL | CG1 | 20.747 | 0.770 | == | | == | !! |
| 74 | VAL | CG2 | 22.071 | 0.947 | == | | == | !! |
| 77 | VAL | CG1 | 22.491 | 0.733 | == | | == | == |
| 77 | VAL | CG2 | 21.784 | 0.755 | == | | | == |
| 81 | LEU | CD1 | 22.548 | 1.077 | == | == | == | == |
| 81 | LEU | CD2 | 26.063 | 1.194 | == | == | == | == |
| 82 | VAL | CG1 | 21.933 | 1.173 | == | | == | == |
| 82 | VAL | CG2 | 21.877 | 1.099 | == | | == | == |
| 87 | VAL | CG1 | 20.901 | 1.033 | (==) | | | !! |
| 87 | VAL | CG2 | 20.479 | 1.151 | (==) | | !! | !! |
| 88 | LEU | CD1 | 25.077 | 0.242 | == | == | == | !! |
| 88 | LEU | CD2 | 20.700 | -0.095 | == | == | == | !! |
| 89 | VAL | CG1 | 21.464 | 0.784 | == | | | !! |
| 89 | VAL | CG2 | 23.461 | 0.711 | == | | == | !! |
| 94 | ILE | CD1 | 13.229 | 0.793 | (==) | == | == | == |
| 106 | LEU | CD1 | 24.657 | 0.440 | == | == | == | == |
| 106 | LEU | CD2 | 23.969 | 0.227 | == | == | == | == |
| 107 | VAL | CG1 | 22.135 | 0.828 | == | | == | !! |
| 107 | VAL | CG2 | 21.775 | 0.745 | == | | | !! |
| 109 | ILE | CD1 | 13.179 | 0.726 | == | == | == | == |
| 112 | LEU | CD1 | 23.297 | 0.652 | == | == | == | == |
| 112 | LEU | CD2 | 26.192 | 0.634 | == | == | == | == |
| 184 | LEU | CD2 | 22.537 | 0.806 | == | == | == | == |
| 190 | VAL | CG1 | 21.380 | -0.176 | == | | == | == |
| 190 | VAL | CG2 | 23.016 | 0.630 | == | | == | == |
| 192 | LEU | CD1 | 24.522 | 1.077 | == | == | == | == |
| 192 | LEU | CD2 | 26.319 | 0.975 | == | == | == | == |
| 194 | ILE | CD1 | 10.361 | 0.506 | == | == | == | == |
| 197 | LEU | CD1 | 23.729 | 0.645 | == | == | == | == |
| 197 | LEU | CD2 | 25.212 | 0.652 | == | == | == | == |
| 201 | LEU | CD1 | 25.766 | 0.784 | == | == | == | == |
| 201 | LEU | CD2 | 22.016 | 0.660 | == | == | == | == |
| 207 | LEU | CD1 | 24.814 | 0.835 | == | == | == | == |
| 207 | LEU | CD2 | 26.319 | 0.762 | == | == | == | == |
| 212 | ILE | CD1 | 14.420 | 0.403 | == | == | == | == |
| 215 | ILE | CD1 | 16.367 | 1.051 | == | == | == | == |
| 217 | VAL | CG1 | 20.710 | 0.916 | == | | | !! |
| 217 | VAL | CG2 | 21.220 | 0.945 | == | | == | !! |
| 223 | ILE | CD1 | 11.794 | 0.782 | == | == | == | == |
| 229 | VAL | CG1 | 21.420 | 1.023 | == | | == | == |
| 229 | VAL | CG2 | 23.133 | 1.180 | == | | == | == |
| 233 | LEU | CD1 | 25.610 | 0.872 | == | == | == | == |
| 233 | LEU | CD2 | 22.819 | 0.696 | == | == | == | == |

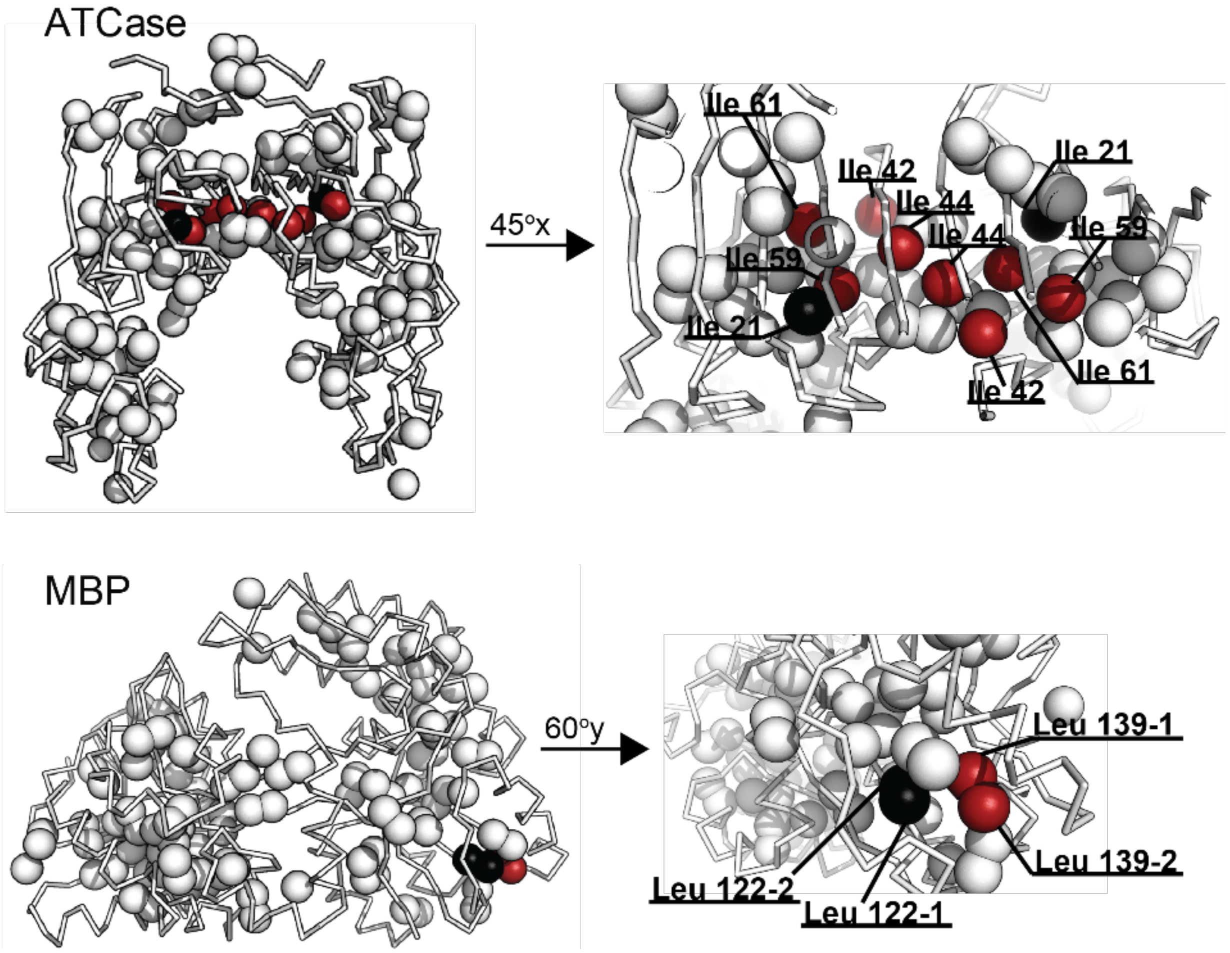

**Figure S4.** Sources of errors in the automatic methyl resonance assignments generated by MethylFLYA. Carbon atoms of the erroneously assigned methyls are shown as red spheres, whereas their correct assignment positions are given in black spheres, or exceptionally in red when the assignment at those positions is also incorrect (i.e. for assignment swaps such as Ile42↔Ile44). The mis-assigned resonances belong to nearby methyl groups. For ATCase, the assignment errors cluster at the interface of the two subunits of the homodimer.

**Table S2.** Summary of errors in the automatic methyl resonance assignments generated by MethylFLYA. The mis-assigned methyls are assigned to spatially proximal residues.

| Protein | Methyl group | Erroneously assigned to | C–C distance (Å) between correct and erroneous assignment |
|---|---|---|---|
| ATCase | Ile 42 $\delta_1$ | Ile 44 $\delta_1$ | 5.5 |
| | Ile 44 $\delta_1$ | Ile 42 $\delta_1$ | 5.5 |
| | Ile 59 $\delta_1$ | Ile 21 $\delta_1$ | 4.0 |
| | Ile 61 $\delta_1$ | Ile 59 $\delta_1$ | 5.9 |
| MBP | Leu 139 $\delta_1$ | Leu 122 $\delta_1$ | 5.2 |
| | Leu 139 $\delta_2$ | Leu 122 $\delta_2$ | 6.9 |

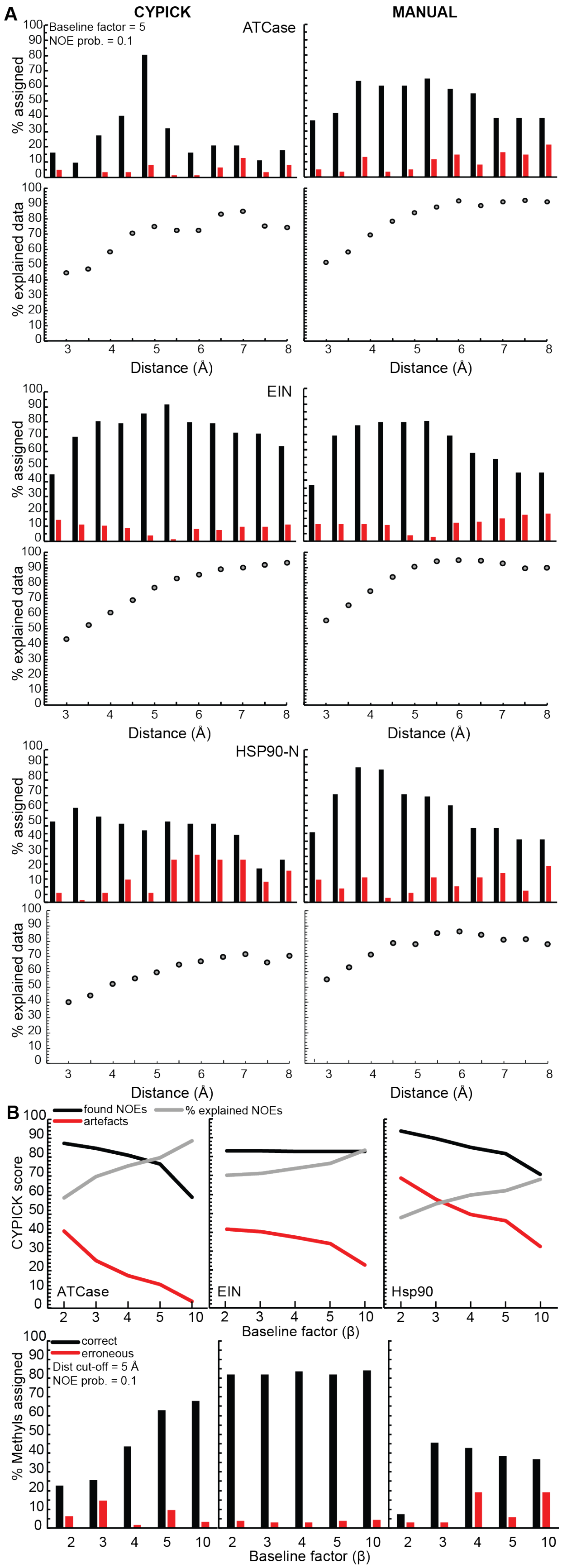

**Figure S5.** Parameter optimization for automatic NOESY peak picking with CYPICK. **(A)** Percentage of accurately (black) and erroneously (red) assigned methyls as a function of methyl $^1$H–$^1$H NOE distance cutoffs, $d_{cut}$, at fixed NOE probability value $p_{NOE} = 0.1$. Results using the automatically generated CYPICK NOESY lists are in the left columns, and results obtained using the manually prepared NOESY lists are in the right column. (**B)** Varying baseline factors for automatic peak picking of methyl-methyl NOESY spectra using CYPICK at fixed distance cutoff $d_{cut}$ = 5 Å and NOE probability value $p_{NOE}$ = 0.1. Variation of the CYPICK find score (*black*), artifact score (red), and the percentage of explained inter-methyl NOEs (*grey*) as a function of the baseline factor. For all three proteins, a fixed distance cutoff $d_{cut}$ = 5 Å and an NOE observation probability $p_{NOE}$ = 0.1 were used to generate the expected methyl-methyl NOEs. Assignment percentages are relative to the number of reference assignments.

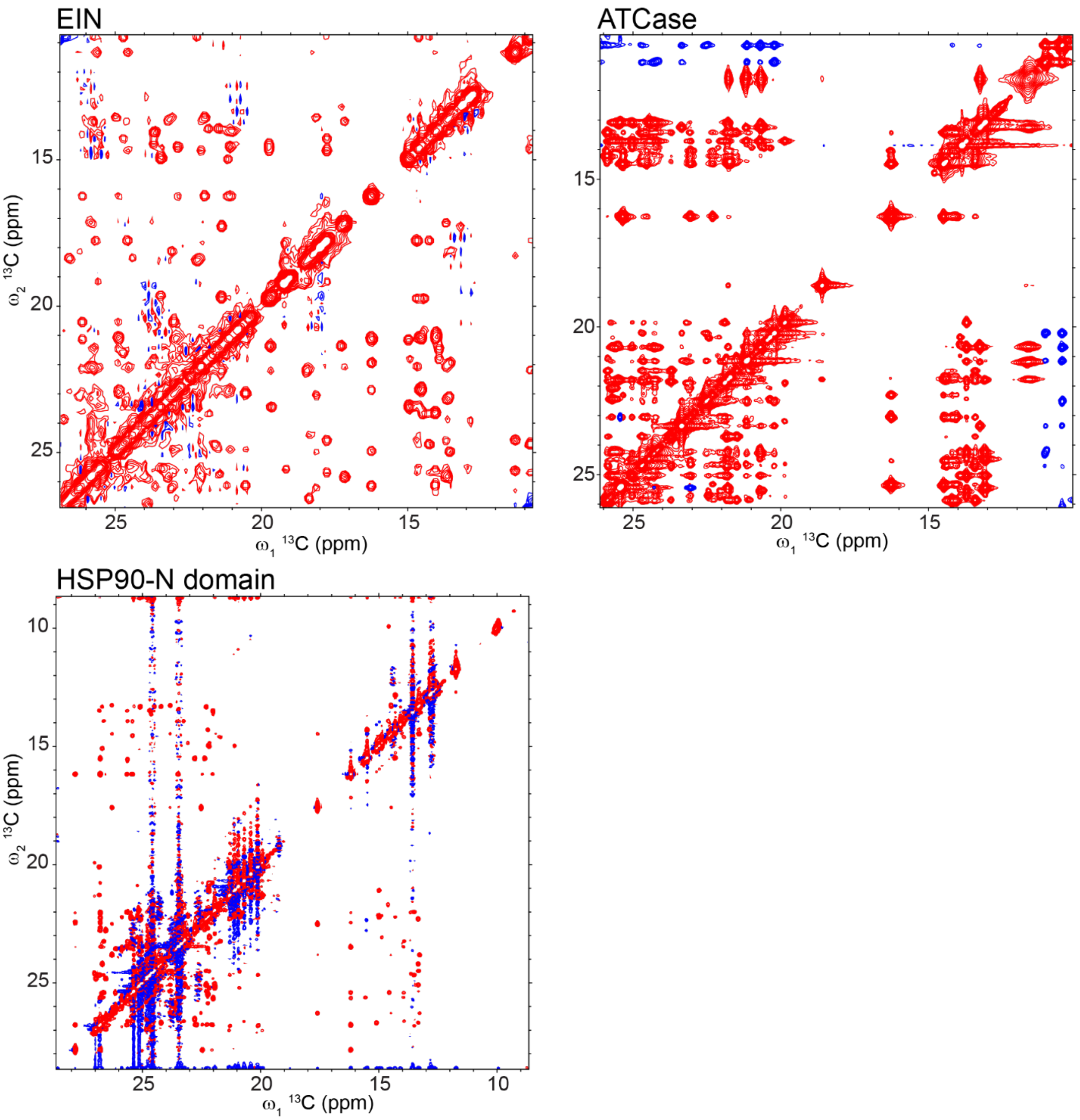

**Figure S6.** 2D $^{13}C(\omega_1)$-$^{13}C(\omega_2)$ projections of 3D CCH NOESY (ATCase, HSP90) or 4D HCCH NOESY spectra (EIN). All spectra are plotted with contour levels starting at a signal-to-noise ratio of three, as determined by the software Sparky (T. D. Goddard and D. G. Kneller, University of California, San Francisco). Positive and negative contours are colored in red and blue, respectively. The spectra were acquired previously by Venditti *et al.*[27] (EIN), Velyvis *et al.*[24] (ATCase), and Shah *et al.*[41] (HSP90).

**Table S3.** MethylFLYA computation times (h) for different combinations of input NMR data, as in Figure 3.

| Protein | Filtered NOEs | Unfiltered NOEs | L, V=LV | 2L, 2V | L, V=LV, 2LV |
|---|---|---|---|---|---|
| EIN | 0.52 | 0.50 | 0.54 | 0.48 | 0.54 |
| ATCase | 0.38 | 0.40 | 0.40 | 0.36 | 0.41 |
| MBP | 0.46 | 0.45 | 0.53 | 0.52 | 0.59 |
| MSG | 1.23 | 1.23 | 1.53 | 1.20 | 1.24 |
| $\alpha_7\alpha_7$ | 0.50 | 0.51 | 0.60 | 0.69 | 0.79 |

Calculations were performed using 100 Intel Xeon E5-2690 processor cores in parallel.

**Table S4.** Results of the CYPICK application to the 3D CCH NOESY (ATCase, HSP90) and 4D HCCH NOESY (EIN) spectra. The results are given for the baseline factor $\beta$ = 5. A find score indicates which percentage of NOE peaks from the reference were also found by CYPICK. Overall, find, and artifact scores are defined by Würz *et al.*[37]

| Protein | Peaks (reference) | Peaks (CYPICK) | Overall score (%) | Find score (%) | Artifact score (%) |
|---|---|---|---|---|---|
| EIN | 618 | 775 | 74 | 83 | 34 |
| ATCase | 563 | 495 | 74 | 77 | 13 |
| HSP90 | 409 | 624 | 68 | 82 | 46 |

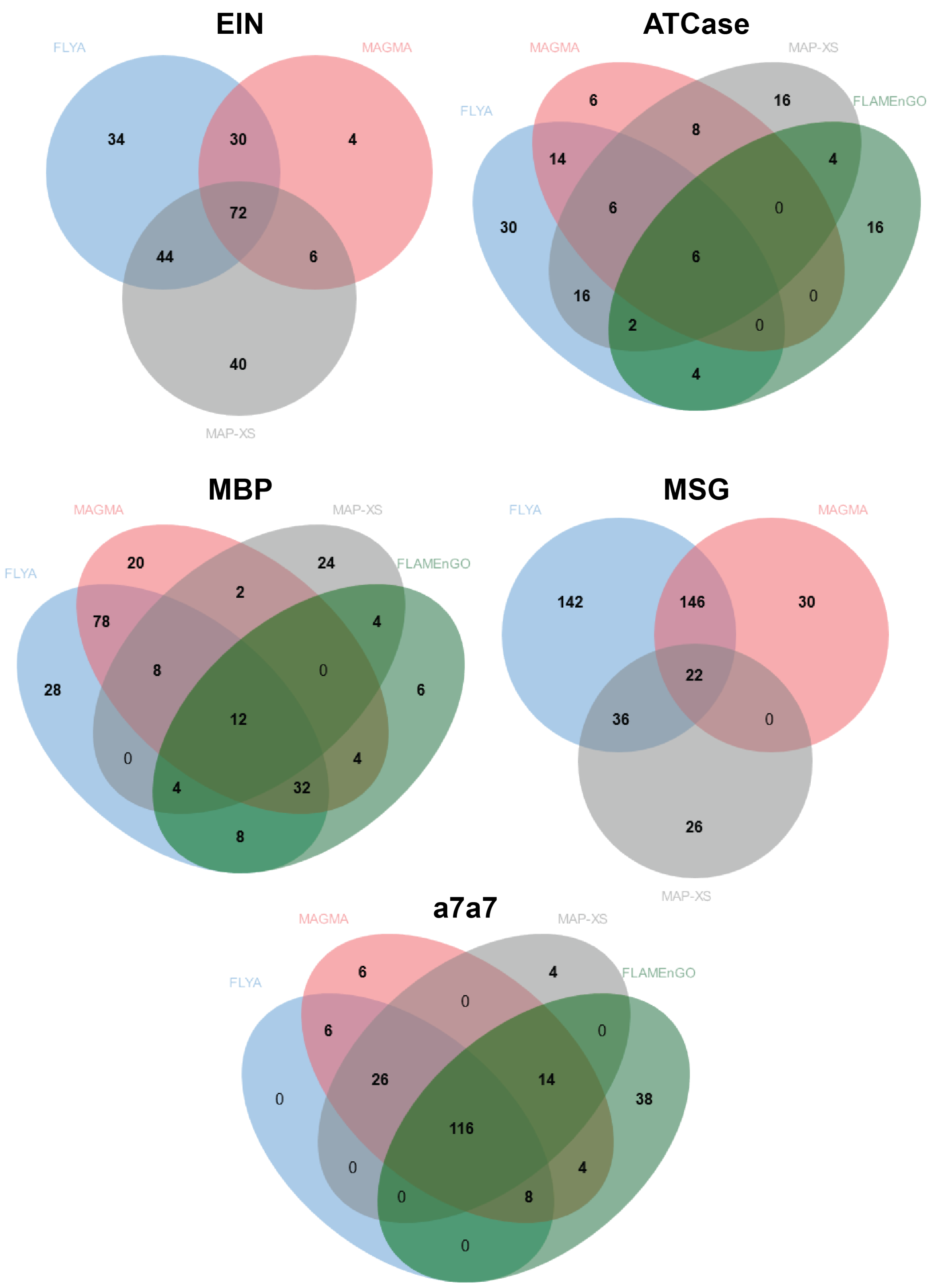

**Figure S7.** Intersection of assignments generated with different automatic methyl assignment protocols. The illustration of the intersections of assignment solutions from MethylFLYA, MAGMA, FLAMEnGO2.0, and MAP-XSII are shown for the indicated benchmark cases. In the case of FLAMEnGO2.0, no confident (100%) assignments were found for EIN and MSG.

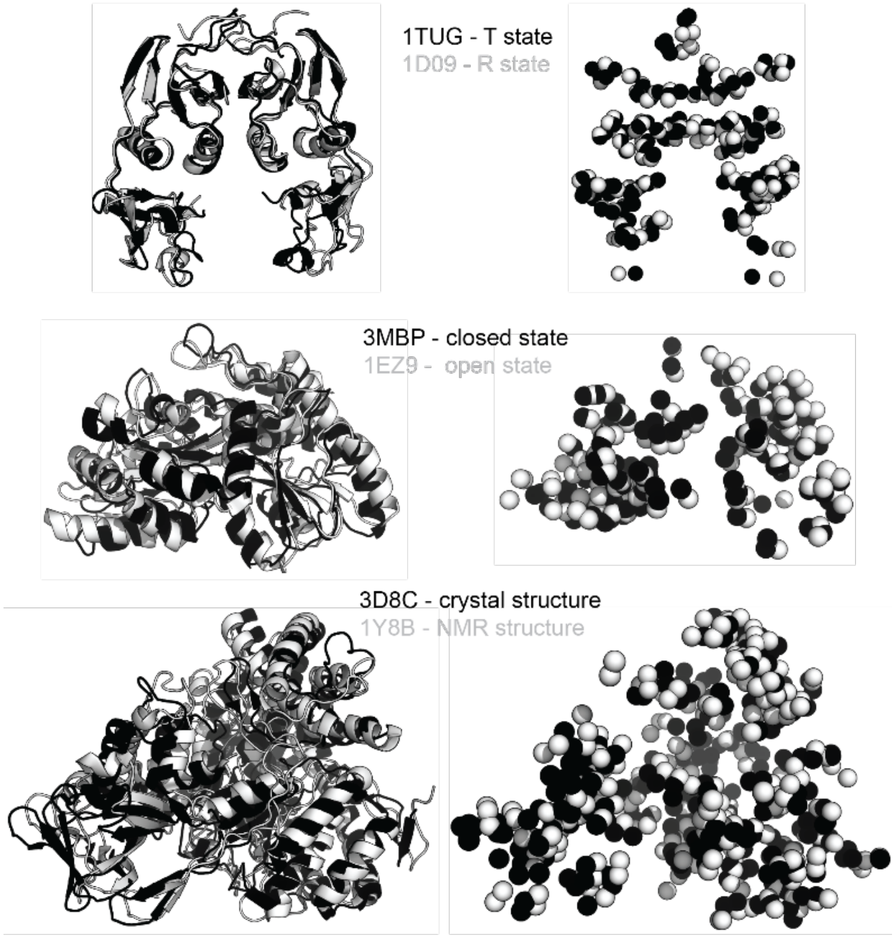

| Protein | PDB ID | correct | error |
|---|---|---|---|
| atcase | 1d09 (Rstate) | 36 | 3 |
| | 1tug (Tstate) | 36 | 1 |
| mbp | 1ez9 (open) | 83 | 2 |
| | 3mbp (closed) | 75 | 2 |
| msg | 1y8b (NMR) | 129 | 0 |
| | 1d8c (X-ray) | 173 | 0 |

**Figure S8.** MethylFLYA performance on different input structures for three enzymes in the benchmark. Differences in backbone conformations between the different protein states are shown with protein structures in cartoon representation (*right* column). In the *left* column, the positions of the methyl carbons are indicated with spheres for each of the conformers, with colors matching those assigned to the backbone (*right*). The total number of accurate and erroneous "strong" methyl assignments generated with MethylFLYA using different conformers is summarized in the table (bottom row).

### Supplementary methods

An example MethylFLYA automated methyl assignment calculation for the N-terminal domain of E. coli Enzyme I (EIN) can be downloaded from http://www.cyana.org/methylflya.tgz.

The complete assignment calculation is performed by first running the *RUN.cya* macro that calls *PREP.cya* to make expected peaks using three different NOE distance cutoffs, and starts parallel FLYA automated assignment runs with *CALC.cya* for the different NOE distance cutoffs. After completing the FLYA runs, consensus chemical shifts are obtained with *CONSOL.cya*, which must be run separately.

Input files:

| | |
|---|---|
| *demo.seq* | amino acid sequence |
| *demo.pdb* | 3D structure |
| *C13HSQC.peaks* | 2D [$^1$H,$^{13}$C]-HMQC peak list with amino acid type assignments (peaks are assigned to methyl groups of the correct amino acid type but arbitrary residue number) |
| *HCcCH.peaks* | short mixing-time 4D CCNOESY for intraresidual Leu/Val connections |
| *CCNOESY.peaks* | 4D CCNOESY peak list, unassigned |
| *ref.prot* | reference chemical shifts (for comparison only) |
| *init.cya* | initialization macro |
| *RUN.cya* | automated assignment calculation (calls *PREP.cya*, *CALC.cya*) |
| *PREP.cya* | prepare peak lists for FLYA |
| *CALC.cya* | run FLYA assignment calculation |
| *CONSOL.cya* | determine consensus chemical shifts |

The preparation macro, *PREP.cya*, performs the following tasks:

1. The [$^1$H,$^{13}$C]-HMQC peak list, *C13HSQC.peaks*, which is assigned to methyls of the correct amino acid type (and arbitrary residue numbers that are not used), is split into four amino acid type-specific peak lists, C13HSQC_*X*.peaks, with *X* = A, I, L, V.
2. The peaks in the unassigned short mixing-time 4D CCNOESY peak list, *HCcCH.peaks* (formally treated as an HCCH TOCSY-type experiment), which contains intraresidual connections between the two methyl groups of Leu or Val, are assigned to amino acid types (and irrelevant, arbitrary residue numbers) according to the closest [$^1$H,$^{13}$C]-HMQC peaks, and the peak list is split into two amino acid type-specific peak lists, *HCcCH_L.peaks* and *HCcCH_V.peaks*.
3. The unassigned 4D NOESY peak list, *CCNOESY.peaks*, is treated similarly, and split into 16 amino acid pair type-specific peak lists, *CCNOESY_XY*.peaks, with *X*, *Y* = A, I, L, V.
4. The macro *peaklists.cya* that specifies the generation of expected peaks during the FLYA calculations in *CALC.cya* is written.

The *RUN.cya* macro creates three subdirectories *demo_d4.5*, *demo_d5.0*, and *demo_d5.5* for the three FLYA calculations with different NOE distance cutoffs, $d_{\text{cut}}$ = 4.5, 5.0, 5.5 Å. In each of these three directories, the input files are copied, the *PREP.cya* macro is executed, and three jobs of 100 individual FLYA assignment runs each are started with *CALC.cya*. The individual assignment runs are executed in parallel on different processors, if available.

Subsequently, consensus assignments are generated with the *CONSOL.cya* macro that produces the following main output files:

| | |
|---|---|
| *consol.prot* | consensus chemical shift lists (in XEASY format) |
| *consol-strong.prot* | strong (confident) consensus chemical shifts (in XEASY format) |
| *consol.tab* | table of consensus methyl assignments |
| *consol.pdf* | plot of consensus methyl assignments |

Optionally (not used in this paper), already known, partial methyl assignments can be included in the calculation by specifying their shifts in a chemical shift list file, e.g. *fix.prot*, and adding the line 'shiftassign_fix := fix.prot' to the *CALC.cya* macro.

For more details on specifying the partial assignments, comparing results to the known reference, or other generic FLYA input file requirements, macros, and output files please see: http://www.cyana.org/wiki/index.php/Tutorials.

### Experiment definitions in the CYANA library

The CYANA library (*cyana.lib*) contains definitions of the experiments necessary for automatic methyl resonance assignment with MethylFLYA:

```
SPECTRUM C13HSQC C H
 0.980 C:C_A* H:H_A*

SPECTRUM CCNOESY3D C1 C2 H1
 0.900 C1:C_A* H1:H_A* ~4.0 H_A* C2:C_A*
 0.800 C1:C_A* H1:H_A* ~4.5 H_A* C2:C_A*
 0.700 C1:C_A* H1:H_A* ~5.0 H_A* C2:C_A*
 0.600 C1:C_A* H1:H_A* ~5.5 H_A* C2:C_A*
 0.500 C1:C_A* H1:H_A* ~6.0 H_A* C2:C_A*

SPECTRUM CCNOESY H1 H2 C2 C1
 0.900 C1:C_A* H1:H_A* ~4.0 H2:H_A* C2:C_A*
 0.800 C1:C_A* H1:H_A* ~4.5 H2:H_A* C2:C_A*
 0.700 C1:C_A* H1:H_A* ~5.0 H2:H_A* C2:C_A*
 0.600 C1:C_A* H1:H_A* ~5.5 H2:H_A* C2:C_A*
 0.500 C1:C_A* H1:H_A* ~6.0 H2:H_A* C2:C_A*

SPECTRUM HCcCH H1 H2 C2 C1
 1.000 H1:H_ALI C1:C_ALI C_ALI C2:C_ALI H2:H_ALI
```

The header line of an experiment definition starts with the word SPECTRUM and gives the name of the spectrum type and a list of labels that correspond to the nuclei that constitute the direct and indirect dimensions, one for each spectral dimension.

Subsequent rows specify the magnetization transfer pathways for the experiment. The first number denotes the peak observation probability. It is followed by a linear list of atom types that defines a molecular fragment, in which atoms must be of the given types, e.g. H_ALI for aliphatic hydrogens, H_A* for aliphatic or (for MethylFLYA irrelevant) aromatic hydrogens, C_ALI for aliphatic carbons, etc., as defined in the ATOMTYPES section at the beginning of the CYANA residue library. Atoms must be connected to the next atom in the list either by a covalent bond or, in the instances where a tilde followed by a number is given, by an NOE, i.e. a distance shorter than the given cutoff (in Å) in the 3D structure. An expected peak is generated whenever a molecular fragment matches the covalent structure and, in case of NOEs, the 3D protein structure. The nuclei for which the frequency is measured in the experiment are identified by labels, followed by a colon. There must be as many labels as in the header (SPECTRUM) line, corresponding to the dimensionality of the spectrum.

See http://www.cyana.org/wiki/index.php/Residue_library_file for more details about the CYANA library.

### Input peak list format

Input peak lists must contain a header line starting with #SPECTRUM that specifies the spectrum type and the labels, which must match the corresponding entry in the CYANA library, but may be permuted to indicate the order in which data for the spectral dimensions is given in peak list columns. For instance, a 4D NOESY peak list in XEASY format may start as follows:

```
# Number of dimensions 4
#SPECTRUM CCNOESY H1 C1 H2 C2
 872   0.230  19.382   0.655  13.602 1 U   1.000E+02  0.000E+00 e 0 -  -  -  -
 874  -0.834  17.697   0.655  13.602 1 U   1.000E+02  0.000E+00 e 0 -  -  -  -
 883   0.848  21.207   0.390  11.341 1 U   1.000E+02  0.000E+00 e 0 -  -  -  -
 887   0.924  22.805   0.390  11.341 1 U   1.000E+02  0.000E+00 e 0 -  -  -  -
 894   1.376  25.567   0.390  11.341 1 U   1.000E+02  0.000E+00 e 0 -  -  -  -
 901   0.407  21.750   0.746  16.119 1 U   1.000E+02  0.000E+00 e 0 -  -  -  -
```

Following the header, data for each peak is given on one line: peak number, peak position (ppm; 4 real numbers), "1 U", peak volume, volume error (if known), "e 0", and assignment ('–' if unassigned). Assignments, if present, are given in the form *A.r*, where *A* denotes an atom name and *r* a residue number.

### Table of consensus chemical shifts

The consolidation of the chemical shift assignments from individual assignment runs into consensus chemical shifts is documented in the MethylFLYA output file *consol.tab*, which contains the final methyl assignment results. The beginning of an example *consol.tab* file is given below.

```
    Atom  Residue      Ref   Shift     Dev  Extent  inside   inref
    QD1   ILE    2            0.739           300.0   100.0     0.0  strong
    CD1   ILE    2           16.246           300.0    99.9     0.0  strong
    QD1   ILE    5            0.843           300.0    37.1     0.0
    CD1   ILE    5           12.874           300.0    62.7     0.0
    QD1   LEU    6   0.755    0.756  -0.001   300.0    99.5   100.0  strong=
    QD2   LEU    6   0.891    0.891   0.000   300.0    96.4    97.0  strong=
    CD1   LEU    6  23.060   23.065  -0.005   300.0    98.3    98.3  strong=
    CD2   LEU    6  25.480   25.557  -0.077   300.0    96.8    97.0  strong=
```

The first three columns in the file list the atom type, residue type, and residue number for each assigned atom. When a reference assignment is known, the value of the known reference chemical shift is listed in the fourth column. The consensus chemical shift (i.e. the MethylFLYA result) is given in the fifth ("Shift") column. When applicable, its deviation from the reference assignment is given in the 'Dev' column in ppm. The 'Extent' column refers to the number of individual assignments runs, in which an assignment for the given atom was obtained. Given that a hundred calculations are run at each of three distance cutoffs, the consolidation runs over 300 individual calculations. The next two columns 'inside' and 'inref' respectively specify the percentage of assignments for this atom in the (300) individual runs that agree (within the chemical shift tolerance specified in the init.cya file) with the consensus or reference assignment, respectively. The final column indicates whether an assignment is 'strong' (i.e. confident). For the calculations in this paper, assignments are classified as strong if and only if the percentage in the 'inside' column is 80% or more. When a reference assignment is known, a '=' is appended to the last column if the MethylFLYA assignment match the reference assignment (within the chemical shift tolerance), or '!' if the two assignments differ by more than the chemical shift tolerance.
